## Supplementary material for "Identification and analysis of individuals who deviate from their genetically-predicted phenotype": Supp Tables 1-8

### Supplementary Information

#### Phenotypic criteria for filtering genes catalogued in OMIM and described as causal for syndromes associated with stature

- Short stature must not be attributable merely to failure to thrive and/or specific metabolic disturbances and/or intestinal failure/enteropathy and/or very severe disease (e.g. early lethality or severe neurologic disease

- Short stature must also be either consistent ($\leq -2$ SD in the great majority of patients with data recorded) or present in multiple families/sibships and accompanied by more severe short stature ($-3$ SD) or skeletal dysplasia (not simply poor bone quality/fractures) or brachydactyly or shortened digits (not clinodactyly or syndactyly) or disporportionate short stature or limb shortening (not simply absence of specific bones)

- Tall stature must be consistent ($> +2$ SD in the great majority of patients with data recorded) or accompanied by more severe tall stature ($> +3$ SD) or arachnodactyly

- Brachydactyly must be not just fifth finger involvement, and either consistent (present in the great majority of patients) or accompanied by consistent short stature or other skeletal dysplasias involving the spine or long bones

- Skeletal dysplasias must involve the spine or long bones (not just the skull or face) and must be accompanied by short stature or brachydactyly as a significant feature (not just clinodactyly or syndactyly), or limb or digit shortening

### Supplementary Tables 1-8 of 10

**STable 1. 238 Genes with prior evidence for a causal association with height, filtered on those with evidence of a dominant inheritance relationship**

| | | | | | | | |
|---|---|---|---|---|---|---|---|
| ACAN | CILK1 | EZH2 | HOXA13 | MAGEL2 | PDE4D | RPS17 | SRCAP |
| ACTB | COL10A1 | FAM111A | HOXD13 | MAP2K1 | PDGFRB | RPS19 | SRY |
| ACTG1 | COL11A1 | FAR1 | HRAS | MAP2K2 | PHEX | RPS24 | STAG2 |
| AFF4 | COL11A2 | FBN1 | HSPA9 | MAP3K7 | PIEZO2 | RPS26 | STAT5B |
| AKT2 | COL1A1 | FBN2 | IFITM5 | MATN3 | PIK3R1 | RPS28 | TBL1XR1 |
| ALDH18A1 | COL1A2 | FBXO11 | IGF1R | MBD5 | POGZ | RPS6KA3 | TBX1 |
| ALPL | COL2A1 | FBXW11 | IGF2 | MC4R | POLE | RPS7 | TBX3 |
| AMER1 | COL9A1 | FGFR1 | IHH | MECP2 | POU1F1 | RRAS2 | TBX4 |
| ANKRD11 | COL9A2 | FGFR2 | IKBKG | MIR140 | PPM1D | RUNX2 | TBX5 |
| ANTXR1 | COL9A3 | FGFR3 | JAG1 | MMP13 | PPP1CB | SEMA3E | TBX6 |
| ARCN1 | COMP | FIG4 | JAK1 | MRAS | PPP2R3C | SGMS2 | TCF4 |
| ARHGAP31 | CREBBP | FLNA | KCNJ2 | MSX2 | PPP3CA | SHOC2 | TGFB2 |
| ARID1A | CSNK2A1 | FLNB | KDM3B | MYCN | PRKAR1A | SHOX | TGFB3 |
| ARID1B | CYP11B1 | FN1 | KDM6A | NAA10 | PTCH1 | SHOX | TGFBR1 |
| ARID2 | CYP19A1 | FZD2 | KIF22 | NEK1 | PTDSS1 | SKI | TGFBR2 |
| ATP8B1 | DDR2 | GDF5 | KMT2A | NF1 | PTH1R | SLC25A24 | THRA |
| ATR | DNMT3A | GH1 | KMT2C | NFIX | PTHLH | SLC2A2 | THRB |
| ATRX | DPF2 | GHR | KMT2D | NIPBL | PTPN11 | SMAD4 | TRPS1 |
| BMP2 | DVL1 | GHSR | KRAS | NLRP3 | PUF60 | SMARCA2 | TRPV4 |
| BMPR1B | DVL3 | GLI2 | LBR | NOG | QRICH1 | SMARCA4 | TTC21B |
| BRAF | EBF3 | GLI3 | LHX4 | NOTCH1 | RAI1 | SMARCB1 | USP9X |
| BRCA1 | EBP | GMNN | LMBR1 | NOTCH2 | RBPJ | SMARCE1 | WASHC5 |
| BRCA2 | EFNB1 | GNAS | LMNA | NPR2 | RIT1 | SMC1A | WDR37 |
| BRPF1 | EP300 | GRIN2A | LOX | NRAS | RNF113A | SMC3 | WNT1 |
| BUB1B | ERCC6 | H1-4 | LRP4 | NSD1 | ROR2 | SNRPB | WNT5A |
| CAMK2G | ESR1 | HCCS | LRP5 | OFD1 | RPL11 | SON | ZBTB18 |
| CBL | EVC | HDAC6 | LTBP3 | OTX2 | RPL13 | SOS2 | ZC4H2 |
| CCNQ | EVC2 | HDAC8 | LZTR1 | P4HB | RPL26 | SOST | ZNF148 |
| CDKN1C | EXT1 | HESX1 | MAB21L2 | PAX8 | RPL35A | SOX11 | SOX9 |
| CHD7 | EXT2 | HMGA2 | MAF | PDE3A | RPL5 | | |

Genes with prior evidence for a causal association with height, filtered on those with evidence of a dominant inheritance relationship

**STable 2. Number of individuals who can be defined as deviating from their genetically predicted height varies substantially be definition**

| **Controls** | 109,858 (69.1%) | |
|---|---|---|
| **Method of Defining Deviator Status** | **Number of Individuals short for their polygenic score (% of population)** | **Number of Individuals tall for their polygenic score (% of population)** |
| **Mahalanobis** (0.001) | 150 (0.094%) | 94 (0.059%) |
| **Mahalanobis** ($0.05/n$) | 10 (0.006%) | 0 (0.00%) |
| **Regression** (2 *s.d.*) | 3,693 (2.32%) | 3,623 (2.28%) |
| **Regression** (3 *s.d.*) | 415 (0.261%) | 287 (0.180%) |
| **GRS Centiles** (1.5 *IQR*) | 693 (0.436%) | 523 (0.329 %) |
| **GRS Centiles** (3 *IQR*) | 23 (0.015%) | 2 (0.001%) |
| **GRS Ranks** (0.001) | 340 (0.021%) | 228 (0.0014%) |
| **GRS Ranks** (1/10,000) | 82 (0.052%) | 44 (0.0028%) |

Number of individuals who are defined as deviating from their polygenic score based on three methodologies, split by relatively tall and relatively short. n = 158,951; IQR = Inter-Quartile Range.

**STable 3.** % **Overlap of individuals defined to be shorter than expected varies substantially between methods**

| | **Mahalanobis (0.001)** | **Mahalanobis (0.05/n)** | **Regression (2 s.d.)** | **Regression (3 s.d.)** | **GRS Ranks (0.001)** | **GRS Ranks (1/10000)** |
|---|---|---|---|---|---|---|
| **Regression (2 s.d.)** | 4.06 | 0.271 | - | - | 0.0915 | 0.0219 |
| **Regression (3 s.d.)** | 31.4 | 2.40 | - | - | 0.781 | 0.195 |
| **GRS Centiles (1.5 IQR)** | 17.7 | 1.44 | 18.8 | 50.3 | 0.486 | 0.117 |
| **GRS Centiles (3 IQR)** | 15.3 | 37.5 | 0.623 | 5.54 | 0.206 | 0.981 |
| **Mahalanobis (0.001)** | - | - | - | - | 0.369 | 0.468 |
| **Mahalanobis (0.05/n)** | - | - | - | - | 0.0264 | 0.108 |

% of overlap between the methods used to determine shorter than expected deviators

**STable 4. % Overlap of individuals defined to be taller than expected varies substantially between methods. No individuals were classified as being relatively tall when using a Mahalanbobis-based P-value threshold = 0.05/n**

| | **Mahalanobis (0.001)** | **Regression (2 s.d.)** | **Regression (3 s.d.)** | **GRS Ranks (0.001)** | **GRS Ranks (1/10000)** |
|---|---|---|---|---|---|
| **Regression (2 s.d.)** | 2.59 | - | - | 0.0629 | 0.0121 |
| **Regression (3 s.d.)** | 21.7 | - | - | 0.746 | 0.153 |
| **GRS Centiles (1.5 IQR)** | 14.3 | 14.4 | 51.1 | 0.436 | 0.0841 |
| **GRS Centiles (3 IQR)** | 2.13 | 0.055 | 0.697 | 0.0952 | 0.704 |
| **Mahalanobis (0.001)** | - | - | - | 0.258 | 0.353 |

% of overlap between the methods used to determine taller than expected deviators

**STable 5. Evidence of enrichment remains consistent across different definitions of misalignment amongst individuals who were short relative to their genetically predicted height.**

| | **Mahalanobis (0.001)** | **Mahalanobis (0.05/n)** | **Regression (2 s.d.)** | **Regression (3 s.d.)** | **GRS Centiles (1.5 IQR)** | **GRS Centiles (3 IQR)** | **GRS Ranks (0.001)** | **GRS Ranks (1/10000)** |
|---|---|---|---|---|---|---|---|---|
| **LoF 0.001 Variants** | 0.014 | 1.00 | $1.00 \times 10^{-4}$ | $1.00{\times}10^{-4}$ | $1.00{\times}10^{-4}$ | 0.025 | $9.00{\times}10^{-4}$ | 0.285 |
| **LoF 0.001 Variants (SS)** | $1.00{\times}10^{-4}$ | 1.00 | $1.00{\times}10^{-4}$ | $1.00{\times}10^{-4}$ | $1.00{\times}10^{-4}$ | $1.00{\times}10^{-4}$ | $1{\times}10^{-4}$ | $1 \times 10^{-4}$ |
| **Missense 0.001 Variants** | 0.159 | 0.750 | 0.377 | 0.326 | 0.269 | 0.623 | 0.485 | 0.936 |
| **Missense 0.001 Variants (SS)** | $1.84{\times}10^{-2}$ | $3.43{\times}10^{-2}$ | $1.00{\times}10^{-4}$ | $5.00{\times}10^{-4}$ | $1.00{\times}10^{-4}$ | $9.58{\times}10^{-2}$ | $4.45{\times}10^{-3}$ | $4.45{\times}10^{-3}$ |
| **ICD9/10 and GP** | $1.00{\times}10^{-4}$ | $1.00{\times}10^{-4}$ | $1.99{\times}10^{-4}$ | $9.00{\times}10^{-4}$ | $3.00{\times}10^{-4}$ | $1.00{\times}10^{-4}$ | $2.00{\times}10^{-4}$ | $1.00 \times 10^{-4}$ |
| **Short (10)** | $1.00{\times}10^{-4}$ | $2.94{\times}10^{-3}$ | $1.00{\times}10^{-4}$ | $1.00{\times}10^{-4}$ | $1.00{\times}10^{-4}$ | $1.00{\times}10^{-4}$ | $1.00{\times}10^{-4}$ | $1.00{\times}10^{-4}$ |
| **TDI** | $1.00{\times}10^{-4}$ | 0.506 | $1.00{\times}10^{-4}$ | $1.00{\times}10^{-4}$ | $1.00{\times}10^{-4}$ | $3.00{\times}10^{-3}$ | $1.00{\times}10^{-4}$ | $1.00{\times}10^{-4}$ |
| **SSHR** | $1.00{\times}10^{-4}$ | $1.00{\times}10^{-4}$ | $1.00{\times}10^{-4}$ | $1.00{\times}10^{-4}$ | $1.00{\times}10^{-4}$ | $3.00{\times}10^{-3}$ | $1.00{\times}10^{-4}$ | $8.70{\times}10^{-3}$ |

Empirical P-values for enrichment in individuals who are short relative to their genetically predicted height across all deviator definitions. SS = Short Stature Specific; LoF = Loss of Function; SSHR = Sitting Standing Height Ratio

**STable 6. Evidence of enrichment remains consistent across different definitions of misalignment amongst individuals who were tall relative to their genetically predicted height. No individuals were classified as being relatively tall when using a Mahalanbobis-based P-value threshold = 0.05/n.**

| | **Mahal anobis (0.001)** | **Regression (2 s.d.)** | **Regression (3 s.d.)** | **GRS Centiles (1.5 IQR)** | **GRS Centiles (3 IQR)** | **GRS Ranks (0.001)** | **GRS Ranks (1/10000)** |
|---|---|---|---|---|---|---|---|
| **LoF 0.001 Variants** | 0.152 | 0.763 | 0.017 | 0.471 | 1 | 0.0434 | 0.182 |
| **LoF 0.001 Variants (TS)** | 0.0237 | $5.57\times10^{-4}$ | 0.0705 | $3.87\times10^{-4}$ | 1.00 | 1.00 | 1.00 |
| **Missense 0.001 Variants** | 0.327 | 0.401 | 0.462 | 0.235 | 0.816 | 0.679 | 0.795 |
| **Missense 0.001 Variants (TS)** | 1.00 | 0.427 | 0.425 | 0.457 | 0.623 | 0.766 | 1.00 |
| **ICD9/10 and GP** | 0.477 | 0.815 | 0.132 | 0.135 | 1.00 | 0.248 | 0.818 |
| **Tall (10)** | $1.00\times10^{-4}$ | $1.00\times10^{-4}$ | $1.00\times10^{-3}$ | $1.00\times10^{-4}$ | 0.382 | $1.00\times10^{-4}$ | $1.00\times10^{-4}$ |
| **TDI** | 0.394 | 0.127 | 0.786 | 0.644 | 1.00 | 0.654 | 0.927 |
| **SSHR** | $1.00\times10^{-4}$ | $1.00\times10^{-4}$ | $1.00\times10^{-4}$ | $1.00\times10^{-4}$ | $1.00\times10^{-4}$ | $1.00\times10^{-4}$ | $2.00\times10^{-4}$ |

Empirical P-values for enrichment in individuals who are tall relative to their genetically predicted height across all deviator definitions. TS = Tall Stature Specific; TDI = Townsend Deprivation Index; SSHR = Sitting Standing Height Ratio

**STable 7. Number of individuals who can be defined as deviating from their genetically predicted LDL-C varies substantially by definition**

| **Controls** | 109,858 (69.1%) | |
|---|---|---|
| **Method of Defining Deviator Status** | **Number of Individuals short for their polygenic score (% of population)** | **Number of Individuals tall for their polygenic score (% of population)** |
| **Mahalanobis** (0.001) | 68 (0.05%) | 90 (0.067%) |
| **Mahalanobis** ($0.05/n$) | 1 (0.0007%) | 2 (0.002%) |
| **Regression** (2 *s.d.*) | 3,085 (2.28%) | 3,128 (2.31%) |
| **Regression** (3 *s.d.*) | 287 (0.213%) | 290 (0.215%) |
| **GRS Centiles** (1.5 $IQR$) | 3,191 (2.36%) | 3,211 (2.37 %) |
| **GRS Centiles** (3 $IQR$) | 13 (0.0096%) | 15 (0.011%) |
| **GRS Ranks** (0.001) | 215 (0.16%) | 222 (0.17%) |
| **GRS Ranks** (1/10,000) | 28 (0.207%) | 31 (0.023%) |

Number of individuals who are defined as deviating from their polygenic score based on three methodologies, split by relatively tall and relatively short. n = 158,951; IQR = Inter-Quartile Range.

**STable 8. Q-risk factor definitions**

| Q-Risk Factor | UKB Data Field |
|---|---|
| **Age** | 21022 |
| **Townsend Deprivation Index** | 21022 |
| **Triglycerides** | 30870 |
| **High Density Lipoprotein (HDL)** | 30760 |
| **Systolic Blood Pressure (SBP)** | 4080 |
| **Diastolic Blood Pressure (DBP)** | 4079 |
| **Body Mass Index (BMI** | 21001 |
| **Height** | 50 |
| **Weight** | 21002 |
| **Cigarettes Per Day (Cigs Per Day)** | 2887 |
| **Alcohol Intake Frequency (Alcohol Freq)** | 1558 |

UKB Data Fields used to derived Q-risk measures
